# Evolutionary dynamics of sex chromosomes of paleognathous birds

**DOI:** 10.1101/295089

**Authors:** Luohao Xu, Simon Yung Wa Sin, Phil Grayson, Scott V. Edwards, Timothy B. Sackton

## Abstract

Standard models of sex chromosome evolution propose that recombination suppression leads to the degeneration of the heterogametic chromosome, as is seen for the Y chromosome in mammals and the W chromosome in most birds. Unlike other birds, paleognaths (ratites and tinamous) possess large non-degenerate regions on their sex chromosomes (PARs or pseudoautosomal regions). It remains unclear why these large PARs are retained over more than 100 MY, and how this retention impacts the evolution of sex chromosomes within this system. To address this puzzle, we analysed Z chromosome evolution and gene expression across 12 paleognaths, several of whose genomes have recently been sequenced. We confirm at the genomic level that most paleognaths retain large PARs. As in other birds, we find that all paleognaths have incomplete dosage compensation on the regions of the Z chromosome homologous to degenerated portions of the W (differentiated regions or DRs), but we find no evidence for enrichments of male-biased genes in PARs. We find limited evidence for increased evolutionary rates (faster-Z) either across the chromosome or in DRs for most paleognaths with large PARs, but do recover signals of faster-Z evolution in tinamou species with mostly degenerated W chromosomes, similar to the pattern seen in neognaths. Unexpectedly, in some species PAR-linked genes evolve faster on average than genes on autosomes, suggested by diverse genomic features to be due to reduced efficacy of selection in paleognath PARs. Our analysis shows that paleognath Z chromosomes are atypical at the genomic level, but the evolutionary forces maintaining largely homomorphic sex chromosomes in these species remain elusive.

## Introduction

Sex chromosomes are thought to evolve from autosomes that acquire a sex determination locus (Bull 1983). Subsequent suppression of recombination between the X and Y (or the Z and W) chromosomes leads to the evolutionary degeneration of the sex-limited (Y or W) chromosome (Bergero and Charlesworth 2009; Bachtrog 2013). Theoretical models predict that suppression of recombination will be favored so that the sexually antagonistic alleles that are beneficial in the heterogametic sex can be linked genetically to the sex determination locus (Rice 1987; Ellegren 2011). Recombination suppression leads to the formation of evolutionary strata, which can occur multiple times in the course of sex chromosome evolution (Lahn and Page 1999; Bergero and Charlesworth 2009; Cortez et al. 2014; Zhou et al. 2014; Wright et al. 2016; Xu et al. 2019). Despite differences in their autosomal origins and heterogamety, eutherian mammals and neognathous birds followed similar but independent trajectories of sex chromosome evolution (Graves 2015; Bellott et al. 2017).

Although this model of sex chromosome evolution has a clear theoretical basis, it is inconsistent with empirical patterns in many vertebrate lineages. Henophidian snakes (boas) are thought to have ZW chromosomes that have remained homomorphic for about 100 MY (Vicoso, Emerson, et al. 2013), although a recent study suggests a transition from ZW to XY system may have occurred (Gamble et al. 2017). Many lineages in fish and non-avian reptiles also possess homomorphic sex chromosomes, in most cases because the sex chromosomes appear to be young due to frequent sex chromosome turnover (Bachtrog et al. 2014). In some species of frogs, homomorphic sex chromosomes appear to be maintained by occasional XY recombination in sex-reversed XY females (the ‘fountain of youth’ model), which is possible if recombination suppression is independent of genotype and instead a consequence of phenotypic sex, such that XY females experience normal recombination (Perrin 2009; Dufresnes et al. 2015; Rodrigues et al. 2018).

Paleognathous birds (Paleognathae), which include the paraphyletic and flightless ratites and the monophyletic tinamous, and comprise the sister group to Neognathae (all other extant birds), also retain largely or partially homomorphic sex chromosomes (de Boer 1980; Ansari et al. 1988; Ogawa et al. 1998; Nishida-Umehara et al. 1999; Pigozzi and Solari 1999; Stiglec et al. 2007; Tsuda et al. 2007; Janes et al. 2009; Pigozzi 2011), albeit with some exceptions (Zhou et al. 2014). These species share the same ancestral sex determination locus, *DMRT1*, with all other birds (Bergero and Charlesworth 2009; Yazdi and Ellegren 2014), and do not fit the assumptions of the ‘fountain of youth’ model (viable and fertile ZW males), requiring an alternative explanation for the retention of homomorphic sex chromosomes. Vicoso and colleagues (Vicoso, Kaiser, et al. 2013), studying the emu, suggested that sexual antagonism is resolved by sex-biased expression without recombination suppression, based on an excess of male-biased gene expression in the pseudo-autosomal region. Alternatively, lack of dosage compensation, which in mammals and other species normalizes expression of genes on the hemizygous chromosome between the homogametic and heterogametic sex, could arrest the degeneration of the W chromosome due to selection to maintain dosage-sensitive genes (Adolfsson and Ellegren 2013). Although these hypotheses are compelling, they have only been tested in single-species studies and without high quality genomes. A broader study of paleognathous birds is therefore needed for comprehensive understanding of the unusual evolution of their sex chromosomes.

Degeneration of sex-limited chromosomes (the W or the Y) leads to the homologous chromosome (the Z or the X) becoming hemizygous in the heterogametic sex. Numerous studies have shown that one common consequence of this hemizygosity is that genes on the X or Z chromosome typically evolve faster on average than genes on the autosomes (Charlesworth et al. 1987; Meisel and Connallon 2013).The general pattern of faster-X or faster-Z protein evolution has been observed in many taxa, including *Drosophila* (Charlesworth et al. 1987; Baines et al. 2008; Avila et al. 2014; Charlesworth et al. 2018), birds (Mank et al. 2007; Mank, Nam, et al. 2010), mammals (Torgerson 2003; Lu and Wu 2005; Kousathanas et al. 2014) and moths (Sackton et al. 2014). One primary explanation for faster-X/Z evolution is that recessive beneficial mutations are immediately exposed to selection in the heterogametic sex, leading to more efficient positive selection (Charlesworth et al. 1987; Vicoso and Charlesworth 2006; Mank, Vicoso, et al. 2010). Alternatively, the degeneration of the Y or W chromosomes results in the reduction of the effective population size of the X or Z chromosomes relative to the autosomes (because there are 3 X/Z chromosomes for every 4 autosomes in a diploid population with equal sex ratios). This reduction in the effective population size can increase the rate of fixation of slightly deleterious mutations due to drift (Mank, Vicoso, et al. 2010; Mank, Nam, et al. 2010). In both scenarios, faster evolution of X- or Z-linked genes is expected.

The relative importance of these explanations varies across taxa. In both *Drosophila* and mammals, faster evolutionary rates of X-linked genes seem to be driven by more efficient positive selection for recessive beneficial alleles in males (Connallon 2007; Meisel and Connallon 2013). However, for young XY chromosomes in plants, reduced efficacy of purifying selection seems to be the cause for the faster-X effect (Krasovec et al. 2018). For female-heterogametic taxa, the evidence is also mixed. In Lepidoptera there is evidence that faster-Z evolution is also driven by positive selection (Sackton et al. 2014) or is absent entirely (Rousselle et al. 2016), whereas in birds, increased fixation of slightly deleterious mutations due to reduced N_e_ is likely a major factor driving faster-Z evolution (Mank, Nam, et al. 2010; Wang et al. 2014; Wright et al. 2015). The non-adaptive effects of faster-Z in birds seem to decrease over time, and the signals of fast-Z effects mostly come from recent non-recombining regions (Wang et al. 2014).

For many paleognaths, a large proportion of the sex chromosomes retains homology and synteny between the Z and the W; these regions are referred to as pseudoautosomal regions (PARs) because they recombine in both sexes and are functionally not hemizygous in the heterogametic sex. In PARs, no effect of dominance on evolutionary rates is expected, and because the population size of the PAR is not different from that of autosomes, an increase in fixations of weakly deleterious mutations is also not expected. Therefore, neither the positive selection hypothesis nor the genetic drift hypothesis is expected to lead to differential evolutionary rates in the PAR compared to autosomes, although other selective forces such as sexually antagonistic selection may impact evolutionary rates in the PAR (Otto et al. 2011; Charlesworth et al. 2014). Moreover, many paleognaths (mainly tinamous) show intermediate or small PARs, implying multiple evolutionary strata, (Zhou et al. 2014) and providing a good system to study the cause of faster-Z evolution at different time scales.

With numerous new paleognath genomes now available (Zhou et al. 2014; Le Duc et al. 2015; Zhang et al. 2015; Sackton et al. 2019), a re-evaluation of sex chromosome evolution in paleognaths is warranted. Here, we investigate faster-Z evolution, dosage compensation and sex-biased expression, to gain a better understanding of the slow evolution of sex chromosomes in ratites. Surprisingly, we did not find evidence for widespread patterns of faster-Z evolution for most paleognaths with large PARs, even when analyzing only differentiated regions (DRs) that are functionally hemizygous in the heterogametic sex. Instead, in a few species we find limited evidence that PARs tend to evolve faster than autosomes. Indirect evidence from the accumulation of transposable elements and larger introns suggests reduced efficacy of selection in both PARs and DRs, potentially because of lower recombination rates compared to similarly sized autosomes. Based on new and previously published RNA-seq data, we find a strong dosage effect on gene expression, suggesting incomplete dosage compensation as in other birds (Itoh et al. 2010; Adolfsson and Ellegren 2013; Uebbing et al. 2013; Uebbing et al. 2015), but do not recover a previously-reported excess of male-biased expression in the PAR (Vicoso, Kaiser, et al. 2013). Our results suggest that simple models of sex chromosome evolution probably cannot explain evolutionary history of paleognath sex chromosomes.

## Results

### Most paleognaths have large pseudoautosomal regions

To identify Z-linked scaffolds from paleognath genomes, we used nucmer (Kurtz et al. 2004) to first align the published ostrich Z chromosome (Zhang et al. 2015) to assembled emu scaffolds (Sackton et al. 2019), and then aligned additional paleognaths (Fig. 1) to emu. We then ordered and oriented putatively Z-linked scaffolds in non-ostrich assemblies into pseudo-chromosomes using the ostrich Z chromosome as a reference (Fig. S1). Consistent with earlier work (Chapus and Edwards 2009), visualization of pseudo-chromosome alignments (Fig. S1) showed little evidence for inter-chromosomal translocations, as expected based on the high degree of synteny across birds (Ellegren 2010); an apparent 12Mb autosomal translocation onto the ostrich Z chromosome is a likely mis-assembly (Fig. S2). This assembly error has been independently spotted using a new linkage map of ostrich (Yazdi and Ellegren 2018).

**FIG. 1.**
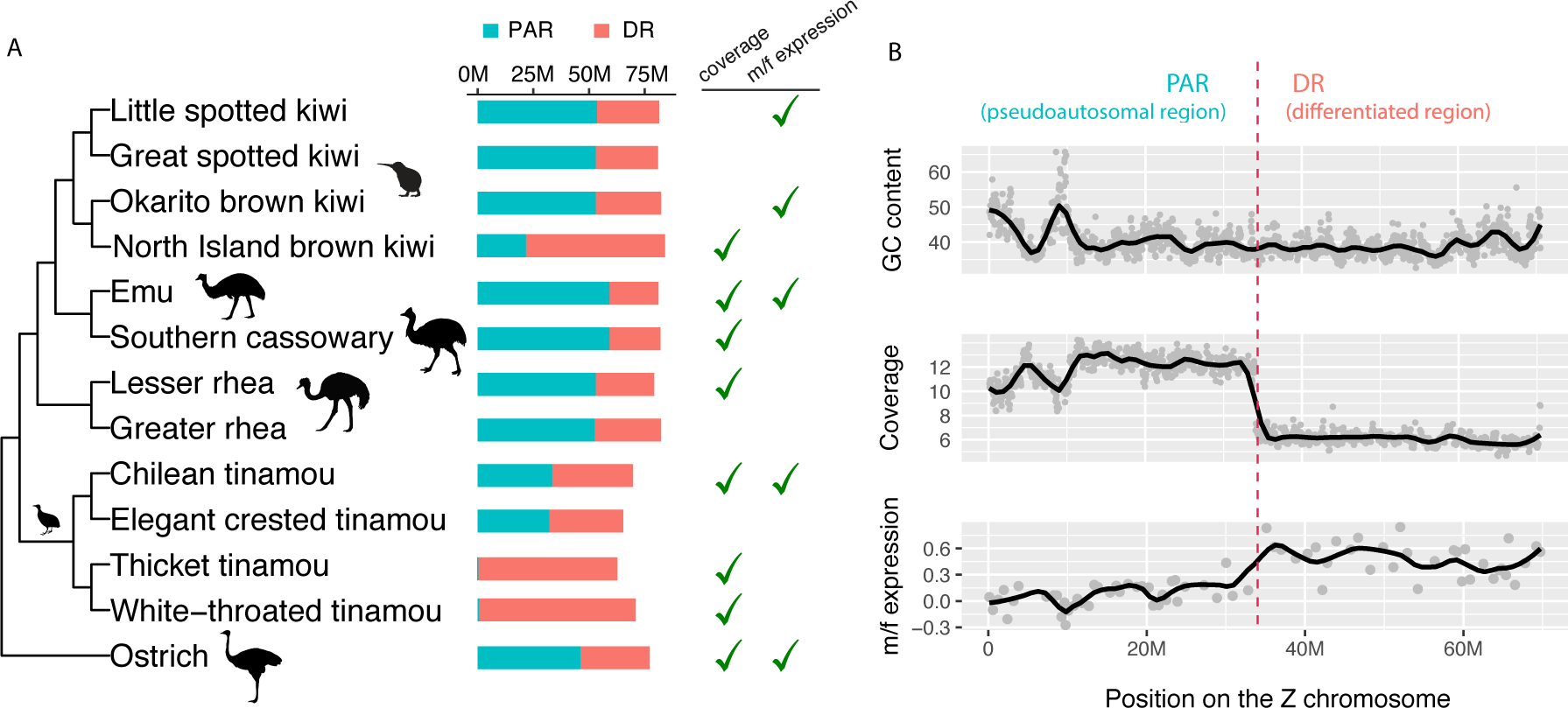
Overview of PAR/DR annotation. **A)** The phylogeny of Palaeognathae based on (Sackton et al. 2019) and (Cloutier et al. 2019). The sizes of the PARs (pseudoautosomal regions) and DRs (differentiated regions) are indicated by the bars in cyan and tomato. The check marks indicate whether the PAR/DR boundaries were annotated by female read coverage and/or male-to-female expression ratios; species with no checks were annotated by homology to closest relatives. **B)** An example of PAR/DR annotation for Chilean tinamou. In the panels of GC content and coverage depth, each dot represents a 50k window. In the panel of m/f expression, each dot represents log2 transformed mean m/f expression ratio of 10 consecutive genes.

We next annotated the pseudoautosomal region (PAR) and differentiated region (DR) of the Z chromosome in each species. In the DR, reads arising from the W in females will not map to the homologous region of the Z (due to sequence divergence associated with W chromosome degeneration), whereas in the PAR, reads from both the Z and the W will map to the Z chromosome. Thus, we expect coverage of sequencing reads mapped to the Z chromosome in the DR to be ½ that of the autosomes or PAR in females, logically similar to the approach used to annotate Y and W chromosomes in other species (Chen et al. 2012; Carvalho and Clark 2013; Tomaszkiewicz et al. 2017). We also annotated PAR/DR boundaries using gene expression data. If we assume that global dosage compensation is absent, as it is in all other birds studied to date (Graves 2014), M/F expression ratios of genes on the Z with degenerated W-linked gametologs in the DR should be larger than that of genes with intact W-linked gametologs in the PAR. There are other processes that can generate a reduced M/F expression ratio in the absence of W chromosome degeneration (e.g., sex-biased expression) and a ‘PAR-like’ M/F expression ratio close to 1 even when the W chromosome is degenerated, such as gene-specific dosage compensation (Naurin et al. 2012) or incomplete degradation of W-linked gametolog. Although these likely account for local departures in expression patterns for individual genes, they are unlikely to explain chromosomal shifts in the means of expression in sliding windows. Nonetheless, we only use expression data when no other method for annotating PAR/DR boundaries is available.

For seven species with DNA (re)sequencing data from females, either newly reported in this study (lesser rhea (*Rhea pennata*), thicket tinamou (*Crypturellus cinnamomeus*), and Chilean tinamou (*Nothoprocta perdicaria*)) or previously published (emu (*Dromaius novaehollandiae*), ostrich (*Struthio camelus*), cassowary (*Casuarius casuarius*), North Island brown kiwi (*Apteryx mantelli*), and white-throated tinamou (*Tinamus guttatus*)), we annotated PAR and DR regions using genomic coverage alone (Fig. 1B, Fig. S3), or in the case of the white-throated tinamou used previously published coverage-based annotations (Zhou et al. 2014). Although some variation in coverage attributable to differences in GC content is apparent, the coverage reduction in the DR region is robust (Fig. 1B). We used expression ratios alone to demarcate the DR/PAR boundaries in little spotted kiwi (*Apteryx owenii*) and Okarito kiwi (*Apteryx owenii*) (Fig. S3), which we found to be in similar genomic locations in both species. For three species (greater rhea (*Rhea americana*), elegant crested tinamou (*Eudromia elegans*) and great spotted kiwi (*Apteryx haastii*)) with neither female sequencing data nor expression data, we projected the DR/PAR boundary from a closely related species (lesser rhea, Chilean tinamou and little spotted kiwi respectively) using shared annotations and synteny.

An alternate approach to identifying the PAR/DR boundary is to rely on SNP densities in females: since the DR is hemizygous in females, we would expect to observe no heterzyogous SNPs in the DR (except for those which arise from mapping of partially degenerated W reads to the Z, which should instead cause an increase in the number of SNPs observed). For most species, SNP data corroborates our PAR/DR boundaries (Fig S3). The exception is the kiwis, where the polymorphism data is ambiguous and suggests the possibility of a recent expansion of the DR and/or a second PAR (Fig S3). We note that the kiwi variation data is based on RNA-seq data from several individuals (Ramstad et al. 2016), and thus it is difficult to rule out biases arising from the interaction between sex chromosome degeneration and transcriptional patterns across the Z chromosome. Thus, we suggest caution in interpreting results from kiwi.

Nonetheless, overall our results corroborate prior cytogenetic studies across paleognaths and support a large PAR in all species except the Tinaminae (thicket tinamou and white-throated tinamou), which have small PARs and heteromorphic sex chromosomes. PAR sizes in large-PAR paleognaths range from ∼20 Mb (23.5% of Z chromosome in North Island brown kiwi) to 59.3 Mb (73% of Z chromosome, in emu); in contrast, PAR sizes in two of the four tinamous and in typical neognaths rarely exceed ∼1 Mb (∼1.3% of Z chromosome size) (Table S1).

### Genes with male-biased expression are not overrepresented in paleognath PARs

Several possible explanations for the maintenance of old, homomorphic sex chromosomes are related to gene dosage (Adolfsson and Ellegren 2013; Vicoso, Kaiser, et al. 2013). We analyzed RNA-seq data from males and females from five paleognath species, including newly collected RNA-seq data from three tissues from emu (brain, gonad, and spleen; 3 biological replicates from each of males and females), as well as previously published RNA-seq data from Chilean tinamou (Sackton et al. 2019), ostrich (Adolfsson and Ellegren 2013), kiwi (Ramstad et al. 2016), and additional embryonic emu samples (Vicoso, Kaiser, et al. 2013). For each species we calculated expression levels for each gene with RSEM (Li and Dewey 2011), and computed male/female ratios with DESeq2 (Love et al. 2014) to assess the extent of dosage compensation, although we note that this measure does not always reflect retention of ancestral sex chromosome expression levels in the hemizygous sex (Gu and Walters 2017). Consistent with previous studies in birds (Graves 2014), we find no evidence for complete dosage compensation by this measure. Instead, we see evidence for partial compensation with M/F ratios ranging from 1.19 to 1.68 (Fig. 2A). The extent of dosage compensation seems to vary among species, but not among tissues within species (Fig. S4). Retention of divergent W-linked gametologs could appear consistent with incomplete dosage compensation, if the reads arising from the W-linked copy no longer map to the Z-linked copy and are thus invisible in the absence of a W assembly. However, previous work in birds suggest that only a very small fraction of Z-linked genes in the DR retain W gametologs (Zhou et al. 2014; Xu et al. 2019), making this explanation unlikely to account for the bulk of expression differences between sexes in the DR.

**FIG. 2.**
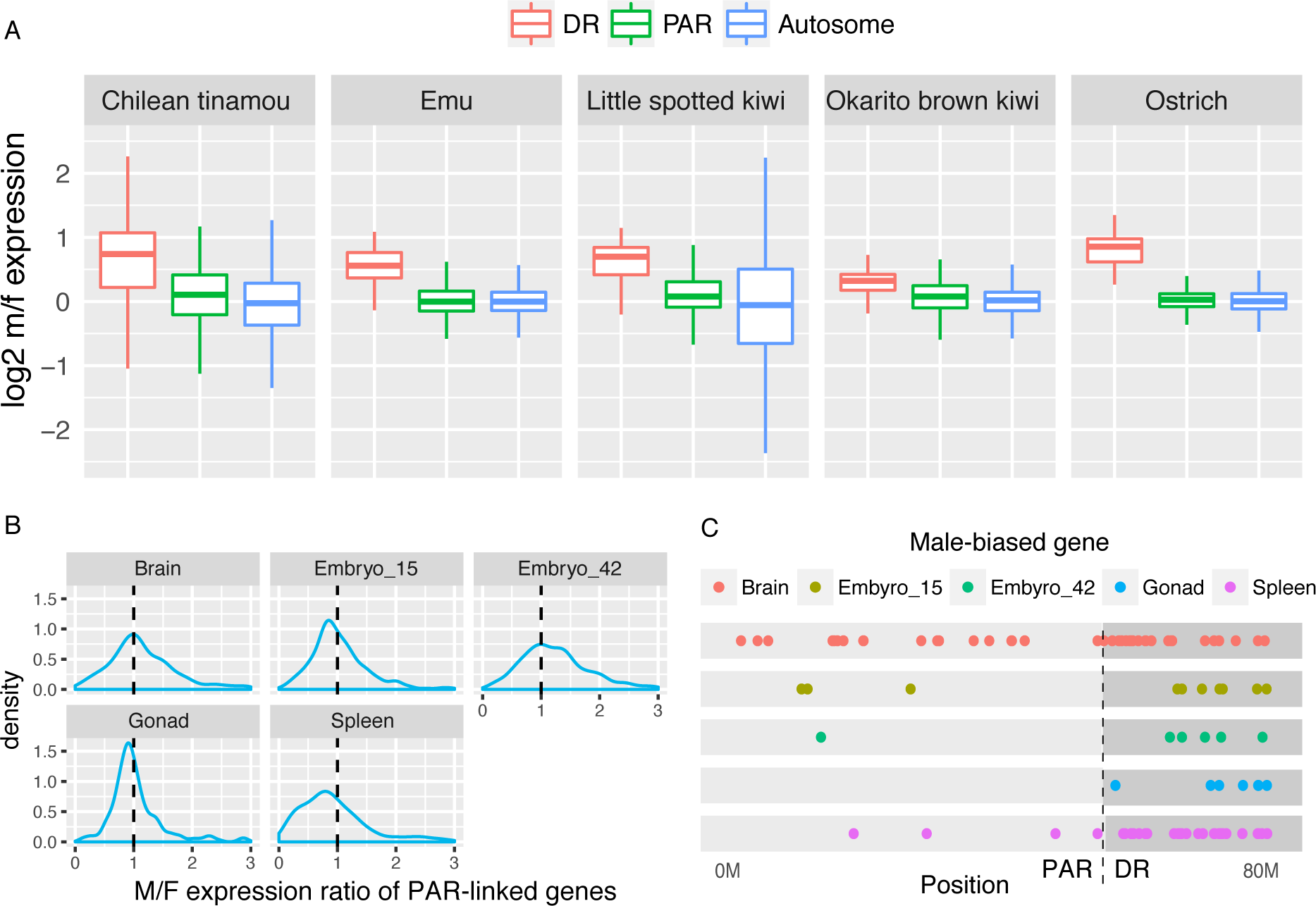
Transcriptomic analyses for five paleognathous species. **A)** Incomplete dosage compensation in emu, kiwi and tinamou. For each species, only one sample is shown: Chilean tinamou (brain), emu (gonad), ostrich (brain) and both kiwis have only blood samples. Log2 m/f expression ratios of DR-linked are larger than 0 but lower than 1. **B)** No excess of male expression levels of PAR-linked genes in most emu tissues, despite slight male-biased expression for 42-day embryo. **C)** No over-representation of male-biased genes in emu PAR. Most Z-linked male-biased genes are located on the DR.

Incomplete dosage compensation poses a challenge for detection of sex-biased genes: higher expression levels of DR-linked genes in males may be due to the incompleteness of dosage compensation rather than sex-biased expression *per se*. With substantially improved genome assemblies and PAR/DR annotations, as well as data from a greater number of species, we re-evaluated the observation that there is an excess of male-biased genes in the emu PAR (Vicoso, Kaiser, et al. 2013). We find that most emu Z-linked male-biased genes are located on the DR (Fig. 2C), and when DR genes are excluded, we no longer detect an excess of male-biased genes on the Z chromosome of emu (p > 0.05 in all tissues, comparing to autosomes, Fisher’s exact test, supplementary Table S2 and Fig. 2C). We similarly do not detect an excess of female-biased genes, either on the Z as a whole or in the PAR only (P > 0.05 in all tissues, Fisher’s exact test, supplementary Table S2). For PAR-linked genes, although there was a slight shift of expression toward male-bias in 42-day old emu embryonic brain (Fig. 2B), only one gene was differentially expressed in male (Fig. 2C). This dearth of genes with male-biased expression in the PAR is largely consistent across other paleognaths with large PARs, including Chilean tinamou, ostrich and little spotted kiwi, with one exception in the Okarito brown kiwi (Fig. S5). Overall, we see little evidence for accumulation of either male- or female-biased genes in paleognath PARs, and suggest that the lack of degeneration of the emu W chromosome and other paleognathous chromosomes is probably not due to resolution of sexual antagonism through acquisition of sex-biased genes.

### Large PARs are associated with lack of faster-Z evolution in paleognaths

The unusually large PARs and the variation in PAR size make Palaeognathae a unique model to study faster-Z evolution. To test whether Z-linked genes evolve faster than autosomal genes, we computed branch-specific dN/dS ratios (the ratio of nonsynonymous substitution rate to synonymous substitution rate) using the PAML free-ratio model for protein coding genes (Yang 2007), based on previously published alignments (Sackton et al. 2019). Because macro-chromosomes and micro-chromosomes differ extensively in the rates of evolution in birds (Gossmann et al. 2014; Zhang et al. 2014) (Fig. S6), we include only the macro-chromosomes (chr1 to chr10) in our comparison, and further focus on only chromosome 4 (97 Mb in chicken) and chromosome 5 (63 Mb) to match the size of the Z chromosome (75 Mb), unless otherwise stated.

We included 23 neognaths and 12 paleognaths in our analysis. Overall, in neognaths, Z-linked genes, with few exceptions, have a significantly higher dN/dS ratio than autosomal (chr 4/5) genes, suggesting faster-Z evolution (Fig. 3). This result is consistent with a previous study involving 46 neognaths (Wang et al. 2014). We further divided Z-linked genes into those with presumed intact W-linked gametologs (PAR genes) and those with degenerated or lost W-linked gametologs (DR genes) to repeat the analysis, because we only expect faster-Z evolution for DR-linked genes. Surprisingly, we do not see widespread evidence for faster-Z evolution in paleognaths for DR genes: only in cassowary, thicket tinamou and white-throated tinamou do DR genes show accelerated dN/dS and dN relative to autosomes (Fig. 4, Fig. S7). Thicket tinamou and white-throated tinamou possess small PARs typical of neognaths, and faster-Z has also been observed for white-throated tinamou in a previous study (Wang et al. 2014), so faster-DR in these species is expected. The observation of faster-DR evolution in cassowary (p = 0.009, two-sided permutation test) suggests that faster-DR evolution may not be limited to species with extensive degeneration of the W chromosome (e.g., with small PARs). However, an important caveat is that the cassowary genome (alone among the large-PAR species) was derived from a female individual, which means that some W-linked sequence could have been assembled with the Z chromosome, especially for the region with recent degeneration. This would cause an artefactual increase in apparent rate of divergence.

**FIG. 3.**
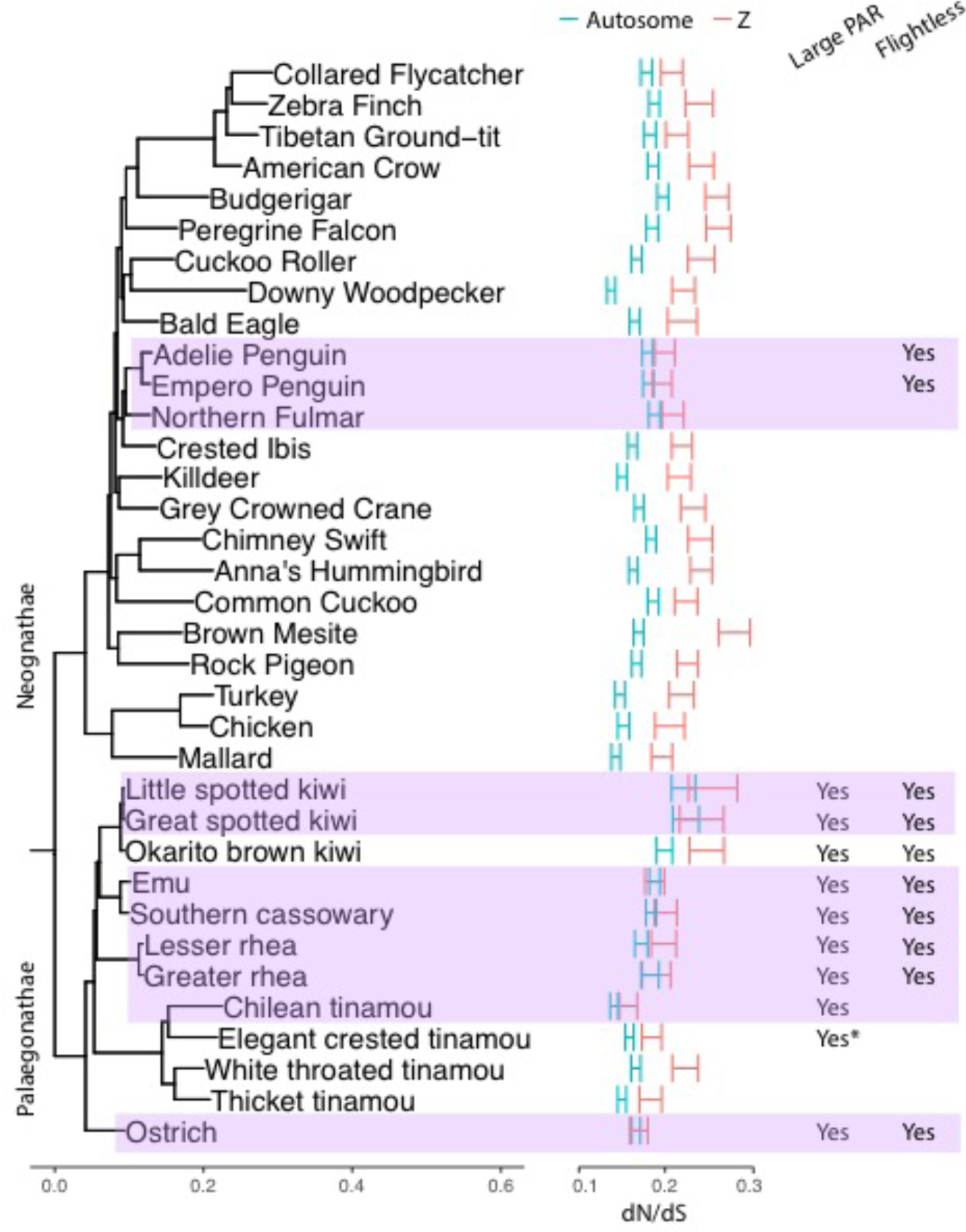
A lack of faster-Z evolution in most Paleognaths. Autosomes were represented by chromosome 4 and chromosome 5 (chr4/5) which have similar sizes compared to the Z chromosomes. The confidence intervals of dN/dS ratios were determined by 1,000 bootstraps. Species without faster-Z effect (permutation test, P > 0.05) are highlighted in purple. The asterisk after ‘Yes’ or ‘No’ indicates uncertainty.

**FIG. 4.**
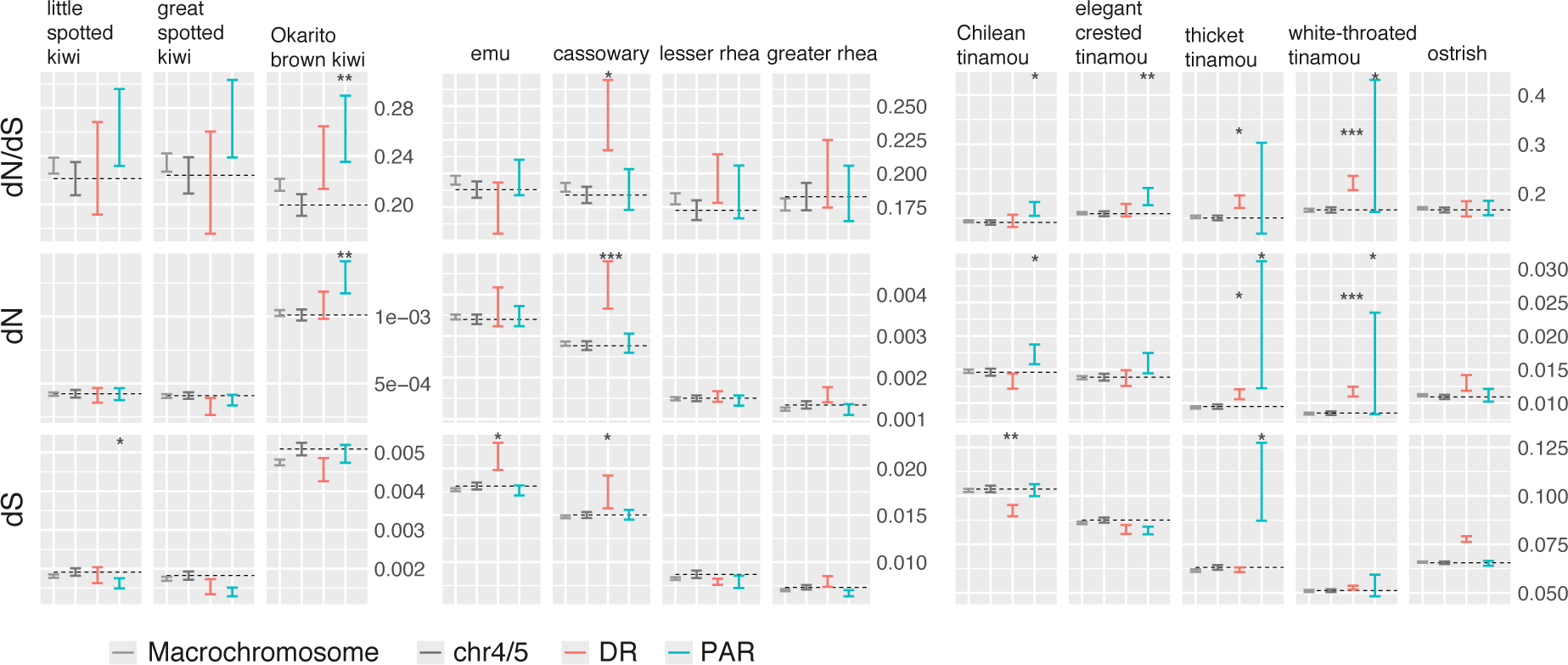
Relative evolutionary rates of Z-linked and autosomal (chr4/5) genes. Confidence intervals were estimated by 1,000 bootstraps. The label chr4/5 stands for chromosome 4 + chromosome 5, and the median value for chr4/5 is also shown as a dotted line. Asterisks indicate the significant levels of PAR/DR *vs.* chr4/5 comparison (two-sided permutation test), * <0.05, ** <0.01, *** <0.001.

Unexpectedly, in three tinamous and one kiwi (white-throated tinamou, Chilean tinamou, elegant-crested tinamou and Okarito brown kiwi) we find evidence that genes in the PAR evolve faster than autosomal genes on chromosomes of similar size (chr4/5), which is not predicted by either the positive selection or genetic drift hypothesis for faster-Z evolution (Fig. 4). All those species have higher dN in the PAR than autosomes, although not significantly so for the elegant-crested tinamou (Fig. 4). Moreover, the faster-PAR effect is not likely to be caused by genes in the newly formed DRs but falsely identified as PARs, because our results are consistent if we remove genes near the inferred PAR/DR boundary (Fig. S8). The faster-PAR in white-throated tinamou is particularly unexpected because previous studies suggest that genes on small PARs evolve slower in birds than non-PAR genes (Smeds et al. 2014). Interestingly, we find the GC content of PAR-linked genes in white-throated tinamou (the only species with both a small PAR and faster-PAR evolution in our analysis) is significantly biased towards GC (Fig. S9), suggesting GC-biased gene conversion might have contributed to the elevated divergence rate. The small number of PAR-linked genes in white-throated tinamou (N=9), however, suggests some caution in interpreting this trend is warranted.

### Evidence for reduced efficacy of selection on the Z chromosome

The signatures of higher dN and dN/dS we observe in the PARs of tinamous and some other species could be driven by increased fixation of weakly deleterious mutations, if the efficacy of selection is reduced in PARs despite homology with the non-degenerated portion of the W chromosome. One potential marker of the efficacy of selection is the density of transposable elements (TEs), which are thought to increase in frequency when the efficacy of selection is reduced (Rizzon et al. 2002; Lockton et al. 2008). We find that chromosome size, which is inversely correlated with recombination rates in birds (Kawakami et al. 2014), shows a strong positive correlation with TE density (lowest in Okarito brown kiwi, r = 0.90; highest in white-throated tinamou, r = 0.98) (Fig. S10, Table S3). Extrapolating from autosomal data, we would expect PARs (smaller than 50Mb in all species) to have lower TE density than chr5 (∼63Mb) or chr4 (∼89Mb) if similar evolutionary forces are acting on them to purge TEs. Strikingly, we find that all paleognaths with large PARs harbor significantly higher TE densities on the PAR than autosomes (Fig. 5), which suggests reduced purging of TEs on PARs.

**FIG. 5.**
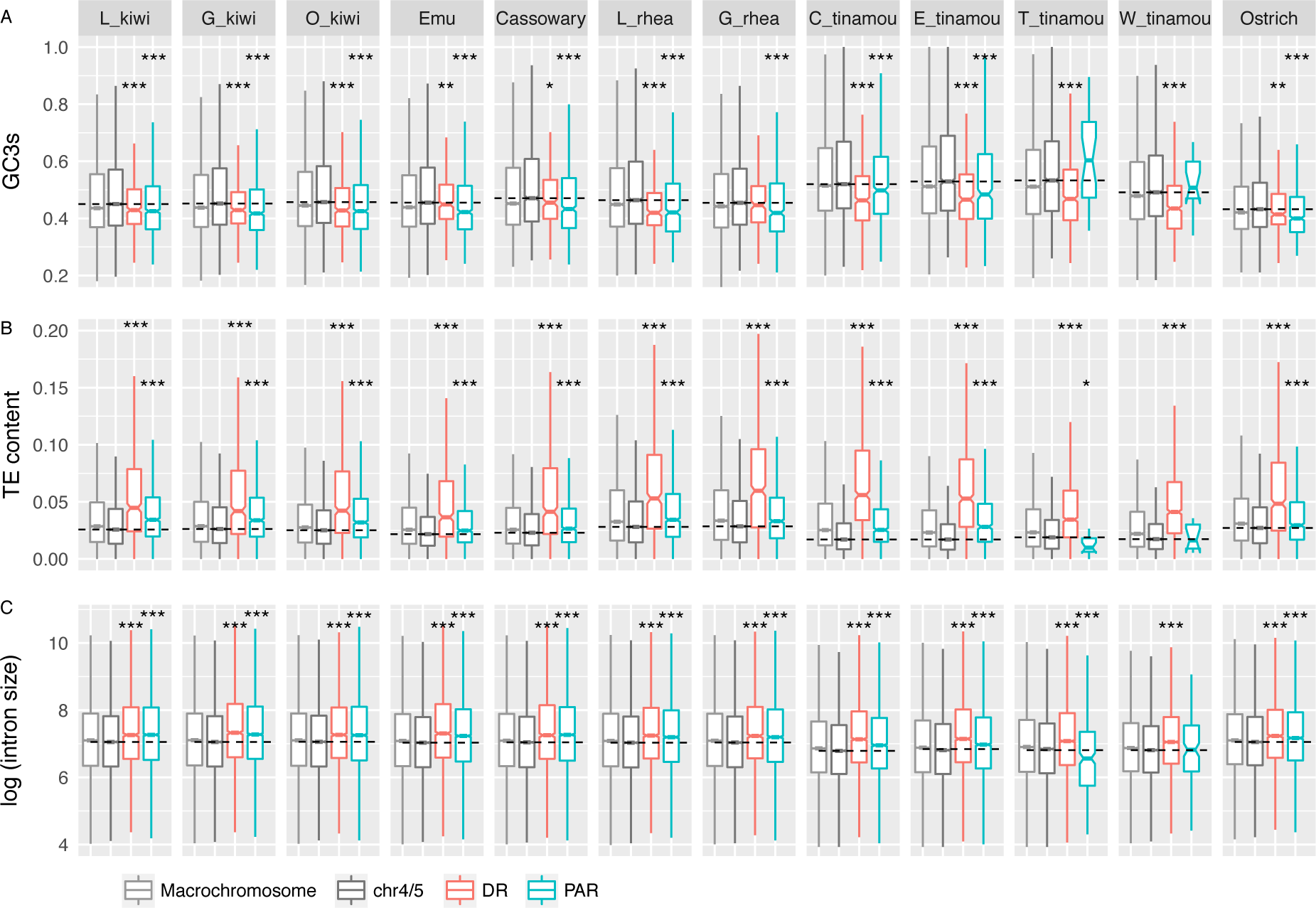
The comparison of PAR/DR *vs.* chr4/5 and macrochromosomes of three genomic features. Median values from chr4/5 are shown as a dotted horizontal line. Asterisks indicate the significant levels of PAR/DR *vs.* chr4/5 comparison (Wilcoxon sum rank test), * <0.05, ** <0.01, *** <0.001 **A)** GC content of the synonymous sites. **B)** TE content, including SINE, LINE, LTR and DNA element. **C)** Log transformed intron size. Abbreviation for species names: L_kiwi, little spotted kiwi; G_kiwi, great spotted kiwi; O_kiwi, Okarito brown kiwi; L_rhea, Lesser rhea; G_rhea, Greater rhea; C_tinamou, Chilean tinamou; E_tinamou, elegant crested tinamou; T_tinamou, thicket tinamou; W_tinamou, white-throated tinamou.

Intron size is probably also under selective constraint (Carvalho and Clark 1999), and in birds, smaller introns are likely favored (Zhang and Edwards 2012; Zhang et al. 2014). If this is also the case in paleognaths, an expansion of intron sizes could suggest reduced efficacy of selection. We compared the intron sizes among PARs, DRs and autosomes across all paleognaths in our study. Like TE densities, intron sizes show strong positive correlation with chromosome size (lowest in Okarito brown kiwi, r = 0.74; highest in thicket tinamou, r = 0.91) (Fig. S10, Table S3). Except for white-throated tinamou and thicket tinamou, intron sizes of the PARs are larger than those of chr4/5 (p < 8.8e-10, Wilcoxon rank sum test, fig. 4C). The pattern of larger intron sizes in the PARs remains unchanged when all macro-chromosomes were included for comparison (Fig. S10). Similar to PARs, DRs also show larger intron sizes relative to chr4/5 (p < 0.00081, Wilcoxon rank sum test).

Finally, codon usage bias is often used as proxy for the efficacy of selection and is predicted to be larger when selection is more efficient (Shields et al. 1988). To assess codon usage bias, we estimated effective number of codon (ENC) values, accounting for local nucleotide composition. ENC is lower when codon bias is stronger, and thus should increase with reduced efficacy of selection. As expected, ENC values showed a strong positive correlation with chromosome sizes (Table S3), and are higher for DR-linked genes in most species (although not rheas, the little spotted kiwi, or the Okarito brown kiwi) (Fig. S11). However, for PAR-linked genes, ENC does not suggest widespread reductions in the efficacy of selection: only cassowary and Chilean tinamou exhibited significantly higher ENC values in the PAR, although a trend of higher ENC values can be seen for most species (Fig. S11).

One possible cause of changes in the efficacy of selection in the absence of W chromosome degeneration is a reduction in the recombination rate of the PAR of some species with a large PAR, although a previous study on the collared flycatcher (a neognath species with a very small PAR) showed that the PAR has a high recombination rate (Smeds et al. 2014). Previous work (Bolivar et al. 2016) has shown that recombination rate is strongly positively correlated with GC content of synonymous third positions in codons (GC3s) in birds, so we used GC3s as a proxy for recombination rate in the absent of pedigree or population samples to estimate the rate directly. We find that GC3s are strongly negatively correlated with chromosome size in all paleognaths (−0.78 ∼ −0.91, p-value <= 0.0068) except for ostrich (r=−0.51, p=0.11) (Fig. S10, Table S3), similar to what was observed in mammals (Romiguier et al. 2010). Recombination rates are also negatively correlated with chromosome sizes in birds (Gossmann et al. 2014; Kawakami et al. 2014) and other organisms (Jensen-Seaman et al. 2004) suggesting that GC3s are at least a plausible proxy for recombination rate. In contrast to the results for collared flycatcher, GC3s of paleognath PARs were significantly lower than those of chr4/5s (p < 2.23e-05, Wilcoxon sum rank test) (Fig. 5A, Fig. S10), except for white-throated tinamou and thicket tinamou. Inclusion of the other macro-chromosome does not change the pattern (p < 0.0034). Moreover, distribution of GC3s along the PAR is more homogeneous compared to chr4 or chr5, except for the 5’prime chromosomal ends (Fig. S12).

## Discussion

Old, homomorphic sex chromosomes have long been an evolutionary puzzle because they defy standard theoretical expectations about how sexually antagonistic selection drives recombination suppression of the Y (or W) chromosome and eventual degradation. The Palaeognathae are a classic example where previous cytogenetic and genomic studies have clearly demonstrated the persistence of largely homomorphic sex chromosomes. Our results extend previous studies, and confirm at the genomic level that all ratites have large, nondegenerate PARs, whereas, in at least some tinamous, degradation of the W chromosome has proceeded, resulting in typically small PARs.

### Evolutionary forces acting on sex chromosomes

Several studies have reported evidence for faster-Z evolution in birds, probably driven largely by increased fixation of weakly deleterious mutations due to reduced N_e_ of the Z chromosome (Mank, Nam, et al. 2010; Wright et al. 2015). However, these studies have focused on neognaths, with fully differentiated sex chromosomes. Here, we show that paleognath sex chromosomes, which mostly maintain large PARs, do not have consistent evidence for faster-Z evolution, although we confirm the pervasive faster-Z effect in neognaths. Notably, the two species in our dataset that presumably share heteromorphic sex chromosomes derived independently from neognaths (white-throated tinamou and thicket tinamou) do show evidence for faster-Z evolution, and in particular faster evolution of DR genes. In contrast, paleognaths with small DR and large PAR do not tend to show evidence for faster-DR, even though hemizygosity effects should be apparent (the exception is cassowary, which may be an artifact due to W-linked sequence assembling as part of the Z).

A previous study on neognaths showed that the increased rate of divergence of the Z is mainly contributed by recent strata, whereas the oldest stratum (S0) does not exhibit the faster-Z effect (Wang et al. 2014). Neognaths and paleognaths share the S0, and, since their divergence, only a small secondary stratum has evolved in paleognaths (Zhou et al. 2014). The absence of a faster-Z effect in paleognath DRs where S0 dominates is therefore largely consistent with the results of the study on the neognath S0. A possible mechanism to explain the lack of faster-Z in the DR is that, in S0, the reduced effective population size (increasing fixation of deleterious mutations) is balanced by the greater efficacy of selection in removing recessive mutations (due to hemizygosity). A recent study on ZW evolution in butterflies suggests a similar model, where purifying selection is acting on the hemizygous DR genes to remove deleterious mutations (Rousselle et al. 2016). Although this model would account for the pattern we observe, it remains unclear why the shared S0 stratum should have a different balance of these forces than the rest of the DR in both neognaths and paleognaths with large DRs. Nonetheless, the evolutionary rates of the DR genes in the older strata are probably the net results of genetic drift and purifying selection against deleterious mutation, with little contribution of positive selection for recessive beneficial mutations.

We also detect evidence for faster evolution of genes in the PAR than for autosomes for three tinamous and one species of kiwi. Because the PAR is functionally homomorphic and recombines with the homologous region of the W chromosome, it is not clear why this effect should be observed in these species. However, a common feature of tinamous and kiwis is that the PARs in some species of these two clades are intermediate or small, e.g. the PARs of North Island brown kiwi and most tinamous. This raises at least two possible explanations for the faster-PAR effect in tinamous and kiwis: 1) the differentiation of the sex chromosomes is more rapid compared with other paleognaths, and at least some parts of the PARs may have recently stopped recombining and actually become DR but undetectable by using the coverage method; or 2) the PARs are still recombining but at lower rate, resulting in weaker efficacy of selection against deleterious mutations. Tinamous are well-known for an increased genome-wide substitution rate compared to other paleognaths (Harshman et al. 2008; Zhang et al. 2014; Zhou et al. 2014; Sackton et al. 2019), but why rates of evolution in the PAR should be so high remains unclear.

### Efficacy of selection and recombination rate

Multiple lines of evidence suggest a possible reduction in the efficacy of selection in the PAR across all paleognaths with a large PAR. Specifically, we find both an increase in TE density and an increase in intron size in PARs. In contrast, we do not find clear evidence for a reduction in the degree of codon bias in PARs. However, it is possible that GC-biased gene conversion (Galtier et al. 2018) and/or mutational bias (Szövényi et al. 2017) may also affect the codon bias, which may weaken the correlation between codon usage bias and the strength of natural selection.

It is unclear, however, why the efficacy of selection may be reduced in PARs. One possible cause is that the PARs may recombine at lower rates than autosomes. This is a somewhat unexpected prediction because in most species PARs have higher recombination rates than autosomes (Otto et al. 2011). In birds, direct estimates of recombination rates of the PARs are available in both collared flycatcher and zebra finch, and in both species PARs recombines at much higher rates than most macrochromosomes (Smeds et al. 2014; Singhal et al. 2015). This is probably due to the need for at least one obligate crossover in female meiosis, combined with the small size of the PAR in both collared flycatcher and zebra finch.

In paleognaths where PARs are much larger, direct estimates of recombination rate from pedigree or genetic cross data are not available. Our observation that GC3s are significantly lower in large paleognath PARs than similarly sized autosomes is at least consistent with reduced recombination rates in these species, although the lower GC3 may alternatively be due to AT mutational bias (Lipinska et al. 2017). A recent study on greater rhea shows that the recombination rate of the PAR does not differ from similarly sized autosomes in females (del Priore and Pigozzi 2017), but this study did not examine males and it cannot exclude the possibilities that the recombination rate in males is lower. A recent study in ostrich, indeed found that the PAR recombines at much lower rate in males than females (Yazdi and Ellegren 2018). If this pattern held true for greater rhea, the sex-average recombination rate of the PAR could potentially be lower relative to similarly sized autosomes. A previous study of emu conducted prior to the availability of an emu genome assembly suggested that the PAR has a higher population recombination rate than autosomes (Janes et al. 2009). However, of twenty-two loci in that study, seven appear to be incorrectly assigned to the sex chromosomes based on alignment to the emu genome assembly (Table S4), potentially complicating that conclusion. The relatively small size of that study and recently improved resources and refined understanding of recombination rates across chromosome types provide opportunities for a new analysis. Further direct tests of recombination rate on ratite Z chromosomes are needed to resolve these discrepancies.

### Sexual antagonism and sex chromosome degeneration

A major motivation for studying paleognath sex chromosomes is that, unusually, many paleognaths seem to maintain old, homomorphic sex chromosomes. We have shown that previously proposed hypotheses do not seem to fully explain the slow degeneration of paleognath sex chromosomes. RNA-seq expression data from both males and females from multiple species suggest dosage compensation is partial in paleognaths, consistent with what has been seen in neognaths. If the absence of complete dosage compensation is the reason for the arrested sex chromosome degeneration in paleognaths, it is not clear why some paleognaths (thicket tinamou and white-throated tinamou) and all neognaths have degenerated W chromosomes and small PARs. The other hypothesis, derived from a previous study on emu (Vicoso, Kaiser, et al. 2013), implies an excess of male-biased genes on the PAR as resolution of sexual antagonism. However, gene expression data from multiple tissues and stages of emu in this study show that male-biased genes are only enriched on the DR, presumably attributable to incomplete dosage compensation and with very few such genes on the PAR. We find similar patterns in other species.

Classic views on the evolution of sex chromosomes argue that recombination suppression ultimately leads to the complete degeneration of the sex-limited chromosomes (Charlesworth et al. 2005; Bachtrog 2006). However, recent theoretical work suggests suppression of recombination is not always favored, and may require strong sexually antagonistic selection (Charlesworth et al. 2014) or other conditions (Otto 2014). Thus, there may be conditions which would have driven tight linkage of the sex-determining locus and sex-specific beneficial loci via the suppression of recombination in neognaths (Gorelick et al. 2016; Charlesworth 2017), but not in paleognaths, although the exact model that could produce this pattern remains unclear, given that it would require, for example, fewer sexually antagonistic mutations in paleognaths than in neognaths. While theoretically possible, there is little evidence to support such a hypothesis, and indeed some paleognaeths (e.g., rheas) have complex mating systems that are at least consistent with extensive sexual conflict (Handford and Mares 1985).

Alternatively, the suppression of recombination between sex chromosomes may be unrelated to sexually antagonistic selection (Rodrigues et al. 2018), and non-adaptive. Simulations suggest that complete recombination suppression can sometimes be harmful to the heterogametic sex, and sex chromosomes are not favorable locations for sexually antagonistic alleles in many lineages (Cavoto et al. 2017). An alternative evolutionary explanation for loss of recombination in the heterogametic sex is then needed. Perhaps the rapid evolution of the sex-limited chromosome may facilitate the expansion of the non-recombining region on the sex chromosome. For instance, once recombination ceases around the sex-determination locus, the W or Y chromosome rapidly accumulate TEs, particularly LTRs, and the spread of LTRs in the non-recombining region may in turn increase the chance of LTR-mediated chromosomal rearrangements, including inversions, leading to the suppression of recombination between the W and Z (or Y and X). Further definition and study of the W chromosomes of paleognaths and neognaths, including patterns of substitution and divergence across genes and noncoding regions, is needed to elucidate the role the W in the evolution of avian sex chromosomes.

## Methods

### Identification of the Z chromosome, PARs and DRs

The repeat-masked sequence of ostrich Z chromosome (chrZ) (Zhang et al. 2015) was used as a reference to identify the homologous Z-linked scaffolds in recently assembled paleognath genomes (Sackton et al. 2019). We used nucmer program (v3.0) from MUMmer package (Kurtz et al. 2004) to first align the ostrich Z-linked scaffolds to emu genome; an emu scaffold was defined as Z-linked if more than 50% of the sequence was aligned. The Z-linked scaffolds of emu were further used as reference to infer the homologous Z-linked sequences in the other paleognaths because of the more continuous assembly of emu genome and closer phylogenetic relationships, and in these cases 60% coverage of alignment was required. During this process, we found that a ∼12Mb genomic region of ostrich chrZ (scf347, scf179, scf289, scf79, scf816 and a part of scf9) aligned to chicken autosomes. The two breakpoints can be aligned to a single scaffold of lesser rhea (scaffold_0) (Fig. S1), so we checked whether there could be a mis-assembly in ostrich by mapping the 10k and 20k mate-pair reads from ostrich to the ostrich assembly. We inspected the read alignments around the breakpoint and confirmed a likely mis-assembly (Fig. S2). The homologous sequences of this region were subsequently removed from paleognathous Z-linked sequences. When a smaller ostrich scaffold showed discordant orientation and/or order, but its entire sequence was contained within the length of longer scaffolds of other paleognaths (Fig. S1), we manually changed the orientation and/or order of that scaffold for consistency. After correcting the orientations and orders of ostrich scaffolds of chrZ, a second round of nucmer alignment was performed to determine the chromosomal positions for paleognathous Z-linked scaffolds.

One way to infer the boundary between the PAR (pseudoautosomal region) and DR (differentiated region) is to compare the differences in genomic sequencing depth of female DNA. Because the DR does not recombine in females and W-linked DRs will degenerate over time and thus diverge from Z-linked DRs, the depth of sequencing reads from the Z-linked DR is generally expected to be half of that for the PAR or autosomes. This approach was applied to cassowary, whose sequence is derived from a female individual. For emu, female sequencing was available from Vicoso et al. (Vicoso, Kaiser, et al. 2013). To facilitate annotation of the PAR, we generated additional DNA-seq data from a female for each of lesser rhea, Chilean tinamou and thicket tinamou. Default parameters of BWA (v0.7.9) were used to map DNA reads to the repeat-masked genomes with BWA-MEM algorithm (Li 2013), and mapping depth was calculated by SAMtools (v1.2) (Li et al. 2009). A fixed sliding window of 50kb was set to calculate average mapping depths along the scaffolds. Any windows containing less than 5kb were removed. Along the pseudo-Z chromosome, the genomic coverage of female reads is usually either similar to that of autosomes (PAR) or reduced to half relative to autosomes (DR). We designated the PAR/DR boundary as the position where a half-coverage pattern starts to appear. For North Island brown kiwi, however, this boundary is unclear, likely due to relatively low quality of the genome assembly. For this reason, as well as a lack of genome annotation for this species, we did not include this species in analyses of molecular evolution.

Another independent method for annotation of the PAR is based on differences in gene expression between males and females for PAR- and DR-linked genes. Because global dosage compensation is lacking in birds and less than 5% of DR-linked genes have homologous W-linked homologs, most DR-linked genes are expected to have higher expression in males. To reduce the effect of transcriptional noise and sex-biased expression, 20-gene windows were used to calculate the mean male-to-female ratios. Increases in male-to-female expression ratios were used to annotate approximate PAR/DR boundaries. This method was applied to little spotted kiwi, Okarito brown kiwi, emu and Chilean tinamou. Given the small divergence between little spotted kiwi and great spotted kiwi, it is reasonable to infer that the latter should have a similar PAR size. Neither female genomic reads nor RNA-seq reads are available for greater rheas and elegant crested tinamou, so the PAR/DR boundaries of lesser rhea and Chilean tinamou were used to estimate the boundaries respectively.

Because the DR is not expected to show heterozygosity in females, we verified the DR annotation by identifying SNPs derived from female sequencing data. To do so, we used GATK (v3.8) pipeline (HaplotypeCaller) following best practices (DePristo et al. 2011). The variants were filtered using parameters ‘QD < 2.0 || FS > 60.0 || MQRankSum < −12.5 || RedPosRankSum < −8.0 || SOR > 3.0 || MQ < 40.0’ and ‘-window 15 -cluster 2’ of the GATK program VariantFiltration. We only retained variants that were heterozygous (allele frequency between 0.2 and 0.8). To calculate the density of female heterozygous sites, the number of variants was counted for every sliding window of 50kb along Z chromosomes. For little spotted kiwi and Okarito brown kiwi, for which only RNA-seq data was available, we called the variants using a similar GATK pipeline, but instead calculated SNPs densities over exons only.

### Comparison of genomic features

To estimate GC content of synonymous sites of the third position of codons (GC3s), codonW (http://codonw.sourceforge.net) was used with the option ‘-gc3s’. The exon density was calculated by dividing the total length of an exon over a fixed 50kb windows by the window size. Similarly, we summed the lengths of transposable elements (TEs, including LINEs, SINEs, LTRs and DNA transposons) based on RepeatMasker outputs (Kapusta *et al*, personal communication) to calculate density for 50kb windows. Intron sizes were calculated from gene annotations (GFF file) using a custom script. Codon usage bias was quantified by the effective number of codons (ENC) using ENCprime (Novembre 2002). We extracted the intronic sequence of each gene for ENCprime to estimate background nucleotide frequency to further reduce the effect of local GC content on codon usage estimates. Wilcoxon sum rank test were used to assess statistical significance.

### Divergence analyses

Estimates of synonymous and non-synonymous substitutions per site were extracted from PAML (Yang 2007) outputs generated by free-ratio branch models, based on previously produced alignments (Sackton et al. 2019). For a given chromosome, the overall synonymous substitution rate (dS) was calculated as the ratio of the number of synonymous substitutions to the number of synonymous sites over the entire chromosome. Outliers (genes showing greater than 1500 substitutions) were removed prior to calculations. Similarly, the chromosome-wide dN was calculated using the numbers of non-synonymous substitutions and sites over the entire chromosome (this is effectively a length-weighted average of individual gene values). The dN/dS values (ω) were calculated by the ratios of dN to dS values. Confidence intervals for dN, dS and dN/dS were estimated using the R package ‘boot’ with 1000 replicates of bootstrapping. P-values were calculated by taking 1000 permutation tests.

### Gene expression analyses

Three biological replicates of samples from emu brains, gonads and spleens of both adult sexes were collected by Daniel Janes from Songline Emu farm (specimen numbers: Museum of Comparative Zoology, Harvard University Cryo 6597-6608). For Chilean tinamou, RNA samples were collected from brains and gonads of both sexes of adults with one biological replicate (raw data from (Sackton et al. 2019), but re-analyzed here). RNA-seq reads for both sexes of ostrich brain and liver (Adolfsson and Ellegren 2013), emu embryonic brains of two stages (Vicoso, Kaiser, et al. 2013), and blood of little spotted kiwi and Okarito brown kiwi (Ramstad et al. 2016) were downloaded from NCBI SRA.

For the newly generated samples (emu brains, gonads and spleens), RNA extraction was performed using RNeasy Plus Mini kit (Qiagen). The quality of the total RNA was assessed using the RNA Nano kit (Agilent). Poly-A selection was conducted on the total RNA using PrepX PolyA mRNA Isolation Kit (Takara). The mRNA was assessed using the RNA Pico kit (Agilent) and used to make transcriptome libraries using the PrepX RNA-Seq for Illumina Library Kit (Takara). HS DNA kit (Agilent) was used to assess the library quality. The libraries were quantified by performing qPCR (KAPA library quantification kit) and then sequenced on a NextSeq instrument (High Output 150 kit, PE 75 bp reads). Each library was sequenced to a depth of approximately 30M reads. The quality of the RNA-seq data was assessed using FastQC. Error correction was performed using Rcorrector; unfixable reads were removed. Adapters were removed using TrimGalore!. Reads of rRNAs were removed by mapping to the Silva rRNA database.

We used RSEM (v1.2.22) (Li and Dewey 2011) to quantify the gene expression levels. RSEM implemented bowtie2 (v2.2.6) to map the RNA-seq raw reads to transcripts (based on a GTF file for each species). Default parameters were used for bowtie2 mapping and expression quantification in RSEM. Both the reference genomes and annotations are from (Sackton et al. 2019). All but the cassowary reference is derived from a female. The raw reads were used for mapping without trimming. TPM (Transcripts Per Million) on the gene level were used to represent the normalized expression. The expected reads counts rounded from RSEM outputs were used as inputs for DESeq2 (Love et al. 2014) for differential expression analysis between sexes. We used a 5% FDR cutoff to define sex-biased genes.

## Acknowledgements

We thank John Parsch, Jamie Walters, Beatriz Vicoso, Daniel Janes and Qi Zhou for their useful comments, the two anonymous reviewers and Judith Mank for helpful comments on the manuscript, and Dan Janes for collecting emu samples. The computations in this paper were performed on the Odyssey cluster at Harvard University and supported by Harvard University Research Computing. This work was supported by NSF grant DEB-135343/EAR-1355292 to SVE. All raw data newly generated in this study (emu RNA-seq, and DNA-seq from lesser rhea, Chilean tinamou, and thicket tinamou) are available from NCBI at BioProject PRJNA450676.

