## Supplemental Materials for "Evolutionary dynamics of sex chromosomes of paleognathous birds"

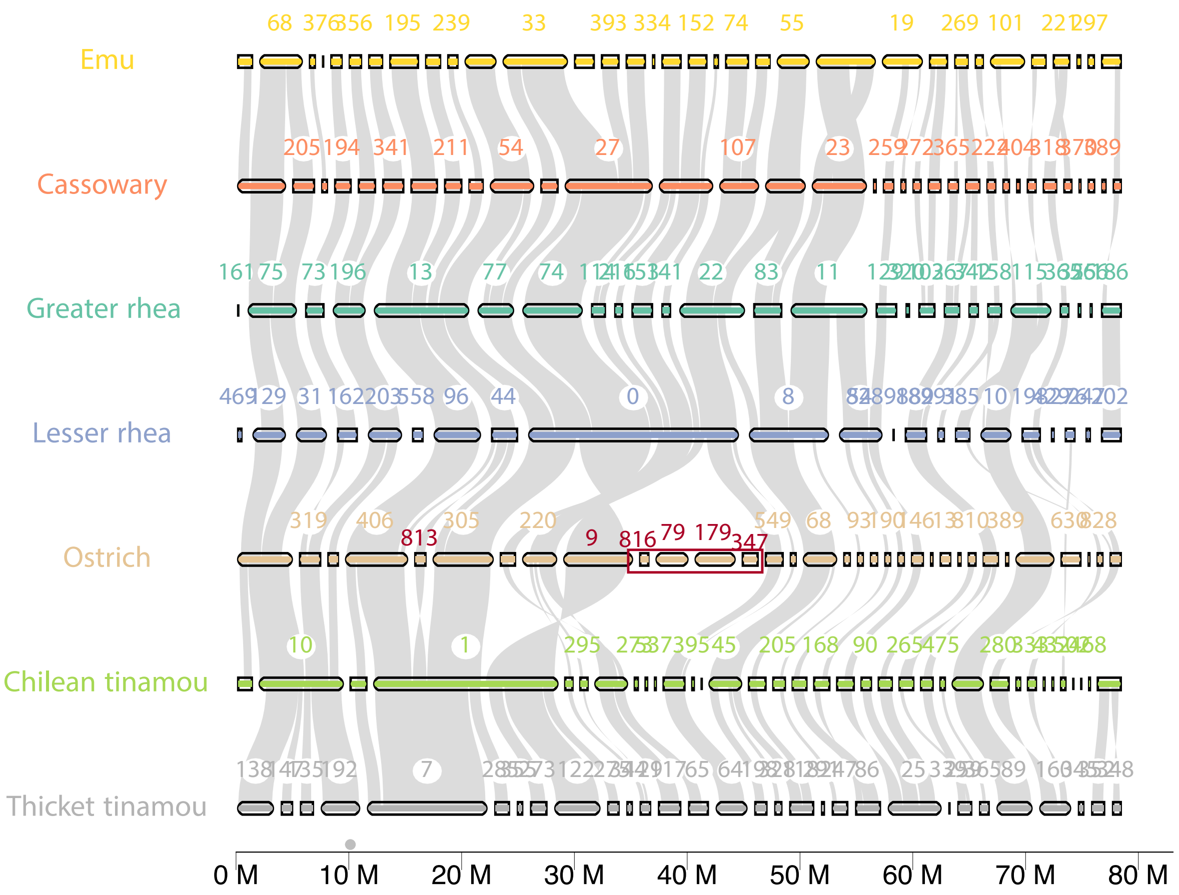


**FIG. S1. Gene synteny among the Z chromosomes**. The alignment of coding sequence and plot were implemented by the python package jcvi (MCscan). Only scaffolds (bars) longer than 50k were shown. As a showcase, the orientation of scaffold 813 of ostrich was corrected. The ~12M containing scaffolds 816, 79, 179, 347 and a part of scaffold 9 were removed from the ostrich. Mate-pair reads alignment for the breakpoint on the scaffold 9 is shown in S2.

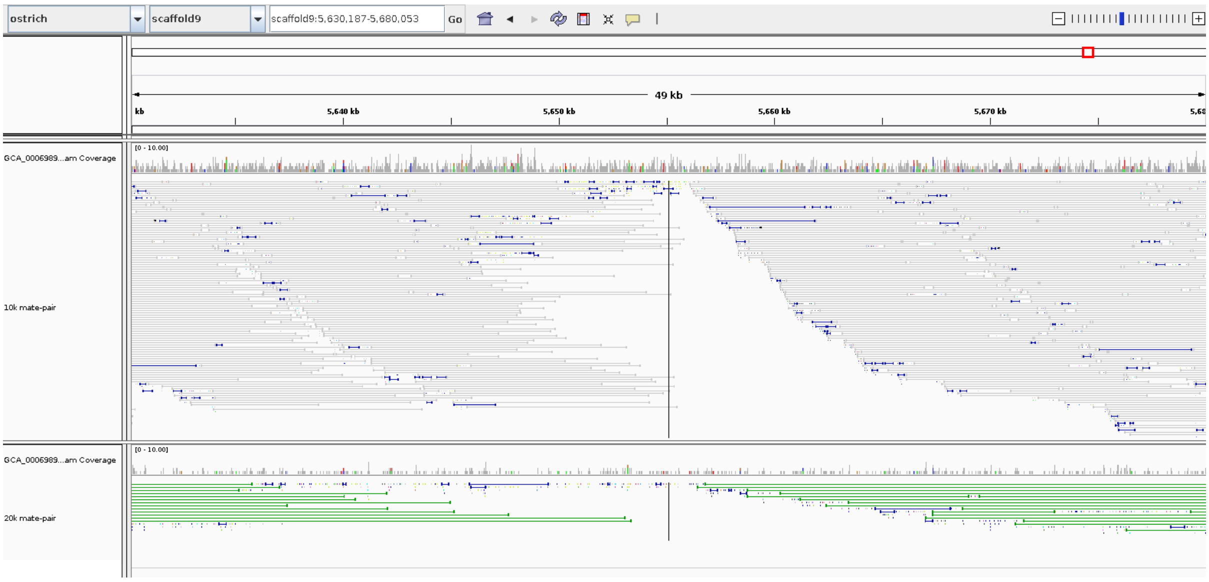


**FIG. S2. Alignments of mate-pair reads against the scaffold9 of ostrich**. The breakpoint is located at the near-end of the scaffold (~5.6M). The upper panel shows the alignments of 10k mate-pair reads and the bottom panel is for 20k mate-pair reads.

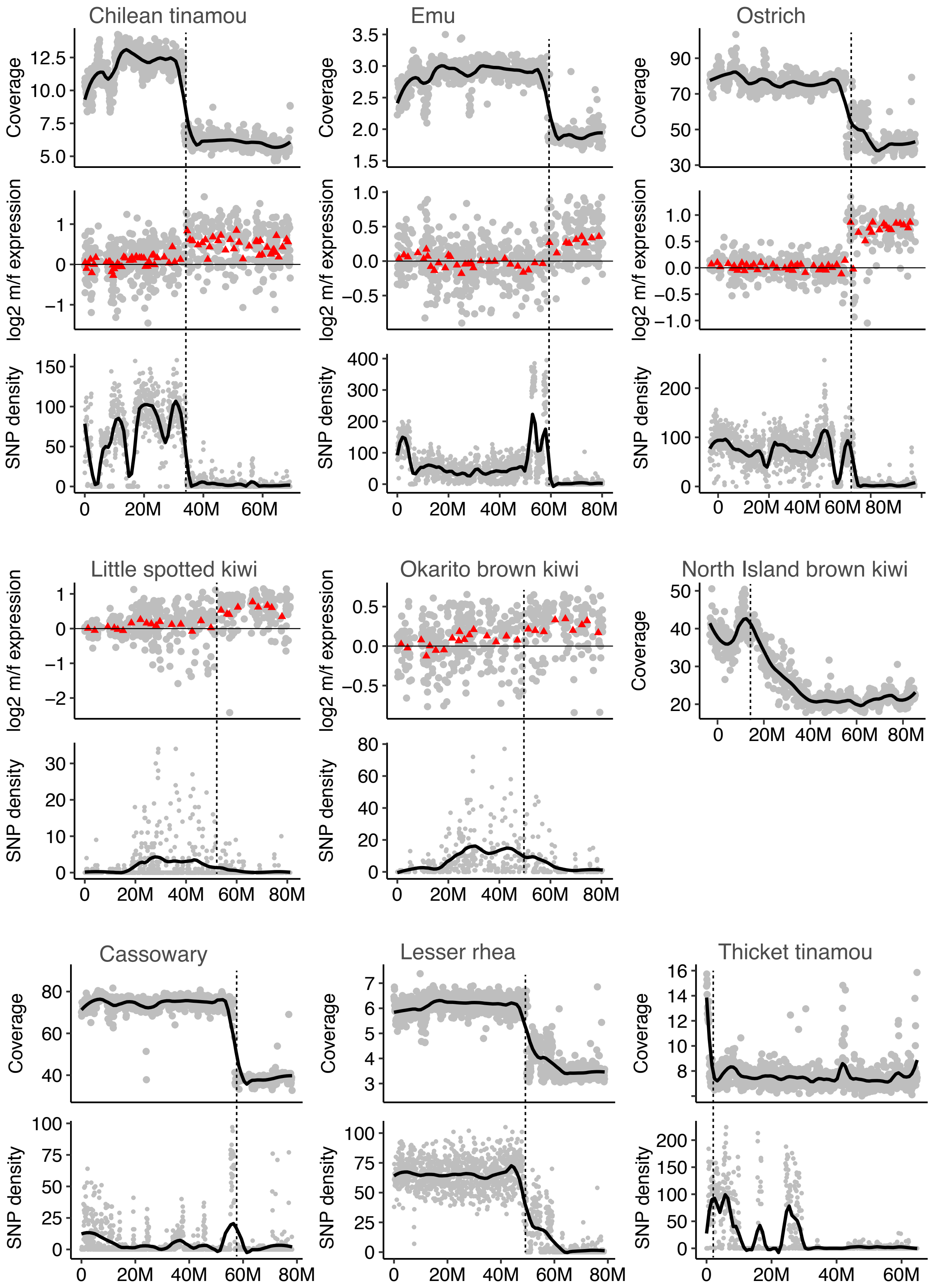


**FIG. S3. Annotation of PAR/DR boundary**. In the 'coverage' panels, each dot represents a 50k window. The black dashed line denotes the boundary of the pseudoautosomal region (PAR) and the differentiated region (DR). In kiwis, an addition dashed line at ~18M show a putative PAR boundary. In 'm/f expression' panels, the red triangle represents the mean m/f expression ratio of 20 genes. The ‘SNP density’ shows the density of female heterozygous sites or SNPs over 50k windows. For both kiwi, female RNA-seq reads were used to call SNPs and the density of SNPs were calculated by dividing the number of SNPs over the length of exonic sequences for every 50k windows. Heterozygous sites were called using GATK pipelines based on female-reads alignments. Note that white-throated tinamou is not shown, as previously published PAR and DR annotations were used for this species.

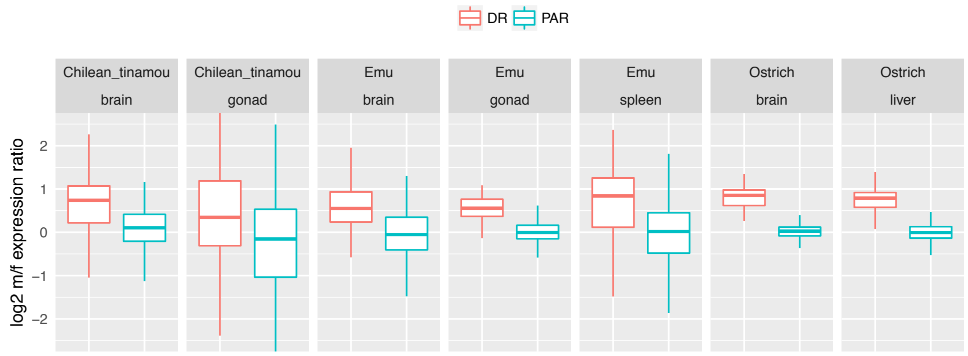


**FIG. S4. Male-to-female expression ratios for DR- and PAR-linked genes**. For Chilean tinamou, emu and ostrich, RNA-seq data of multiple tissues of both sexes are available. The m/f ratios (log2 transformed) of DR-linked genes are larger than 1 but less than 2, suggesting incomplete dosage compensation, but show limited variation within species.

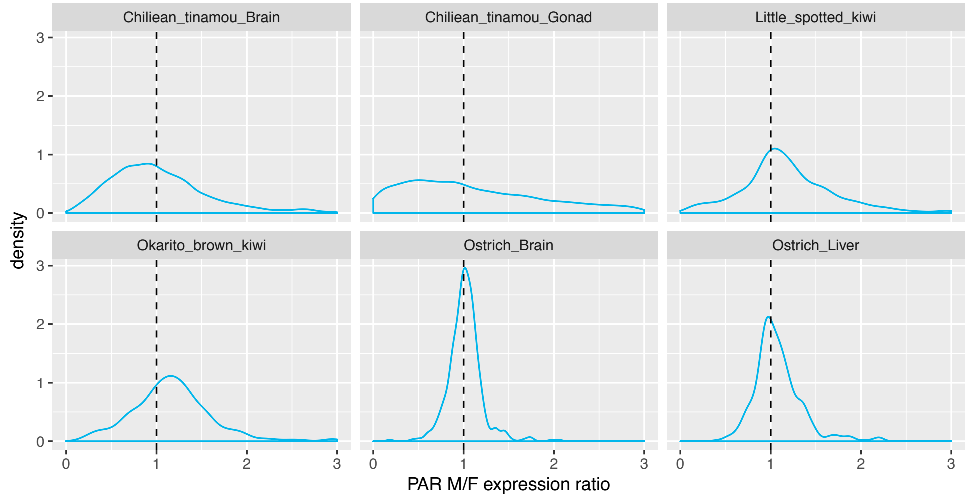


**FIG. S5. Distribution of male-to-female expression ratios for PAR-linked genes**. In most samples m/f expression ratios do most deviate from 1, suggest similar expression levels of PAR-linked genes between males and females. Only in Okarito brow kiwi, however, male expression levels are slightly higher than for females.

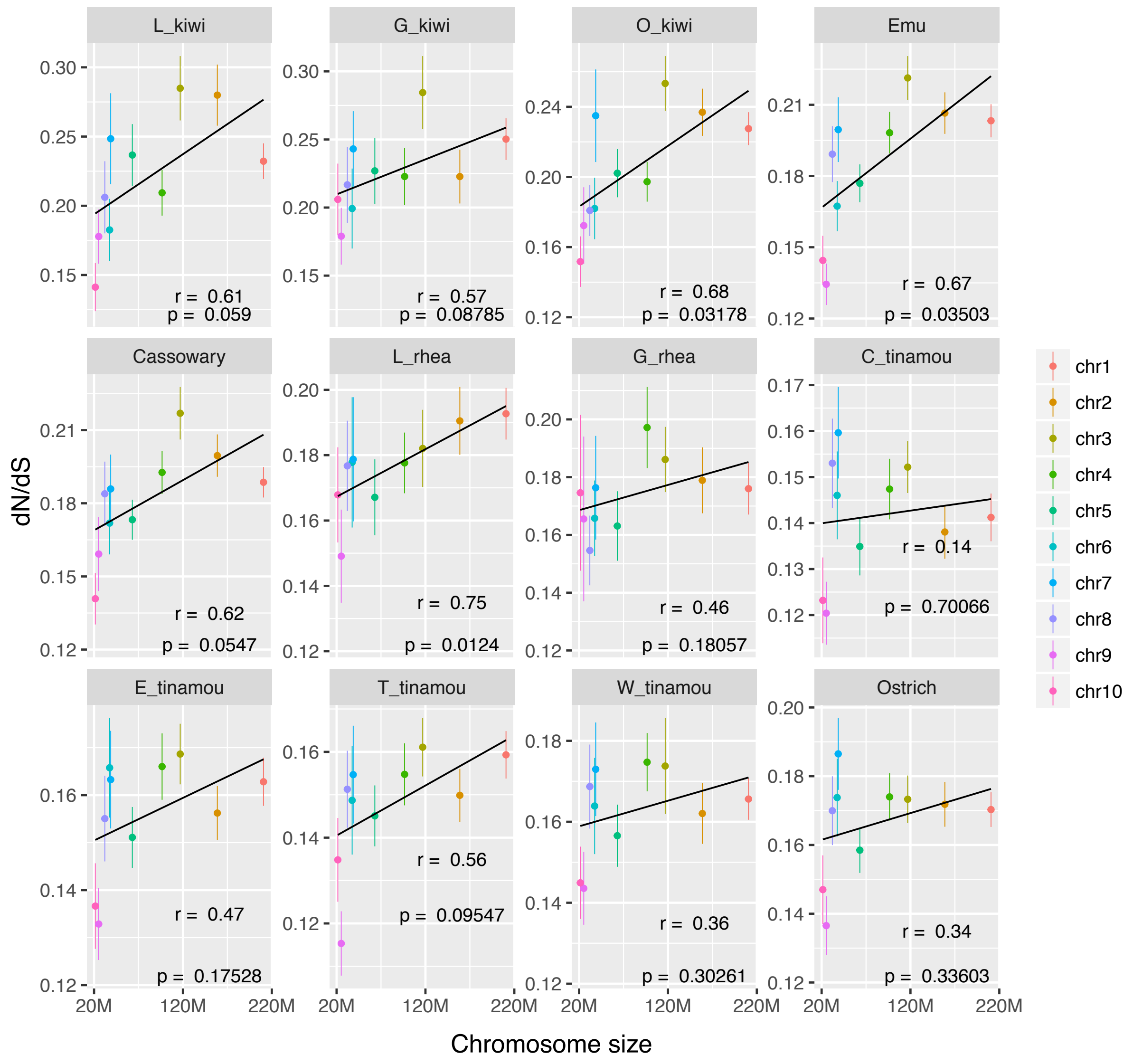


**FIG. S6. Positive correlation of dN/dS ratios and chromosome size among macro-chromosomes**. Among macro-chromosome (chr1 – chr10), chromosome size positively correlates with dN/dS ratios. The chromosome size of the Z is about 75M, between the sizes of chr4 (~97M) and chr5 (~63M). The ‘r’ strands for Pearson’s correlation coefficient. Abbreviation for species names: L_kiwi, little spotted kiwi; G_kiwi, great spotted kiwi; O_kiwi, Okarito brown kiwi; L_rhea, Lesser rhea; G_rhea, Greater rhea; C_tinamou, Chilean tinamou; E_tinamou, elegant crested tinamou; T_tinamou, thicket tinamou; W_tinamou, white-throated tinamou.

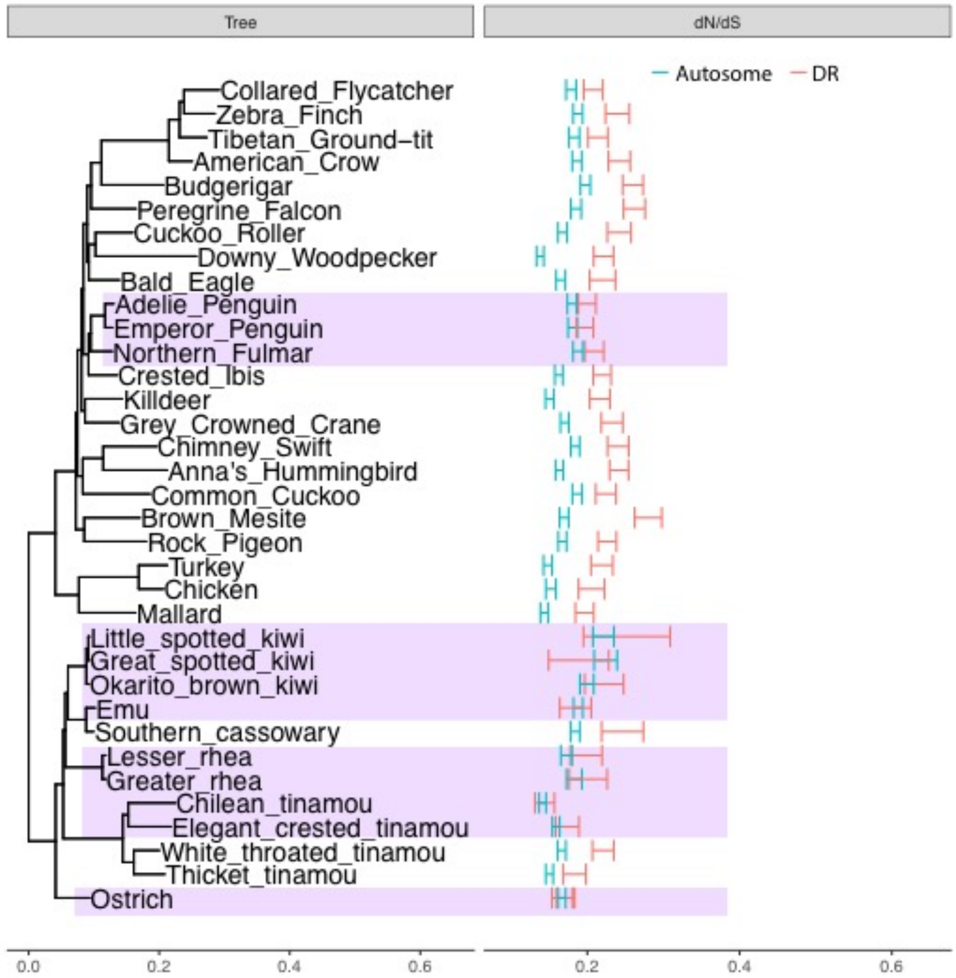


**FIG. S7. A lack of faster-DR in most palaeognaths**. The PAR-linked genes were removed from the analysis. Species without faster-DR effect (permutation test, P > 0.05) were highlighted by purple colour. The faster-Z effect is no longer observed in Okarito brown kiwi, elegant crested tinamou and thicket tinamou after PAR-linked genes were removed.

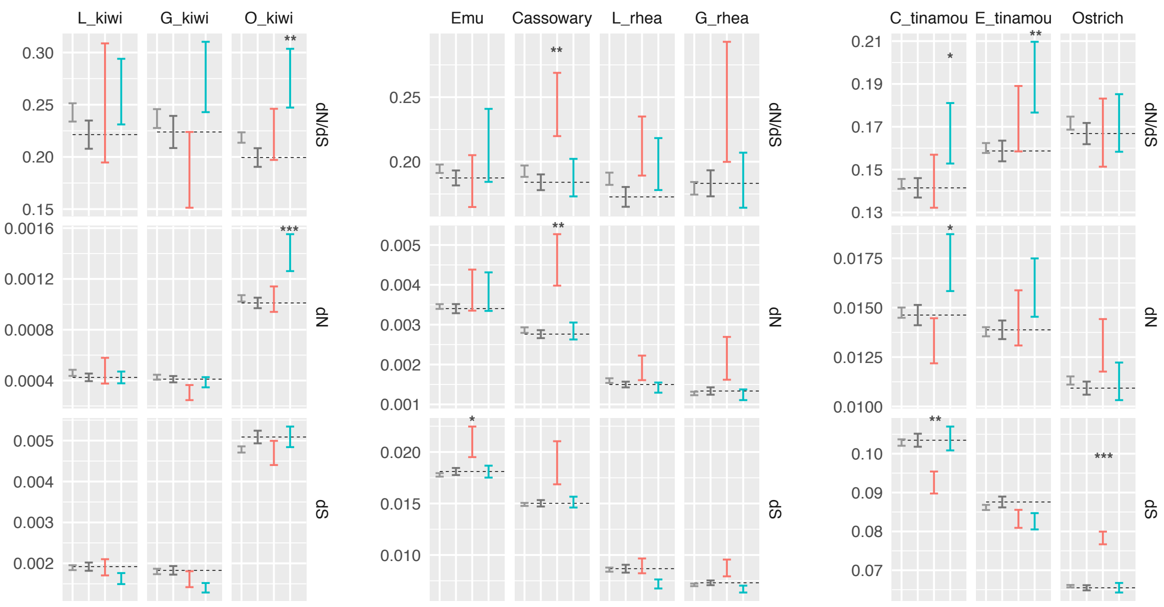


**FIG.S8. The boundary of PAR/DR does not show faster-Z effect**. dN (nonsynonymous substitution rate), dS (synonymous substitution rate) and their ratios (dN/dS) are shown for PAR (cyan), DR (red), chr4/5 (dark grey) and macro-chromosome (grey) genes. The test for faster-Z evolution was repeated after the exclusion of PAR-linked gene close to PAR boundaries (less then 5 Mb away). Similar to Fig. 4 in the main text, Confidence intervals were estimated by 1,000 bootstraps. Asterisks indicate the significant levels of PAR/DR *vs.* chr4/5 comparison (two-sided permutation test), * <0.05, ** <0.01, *** <0.001.

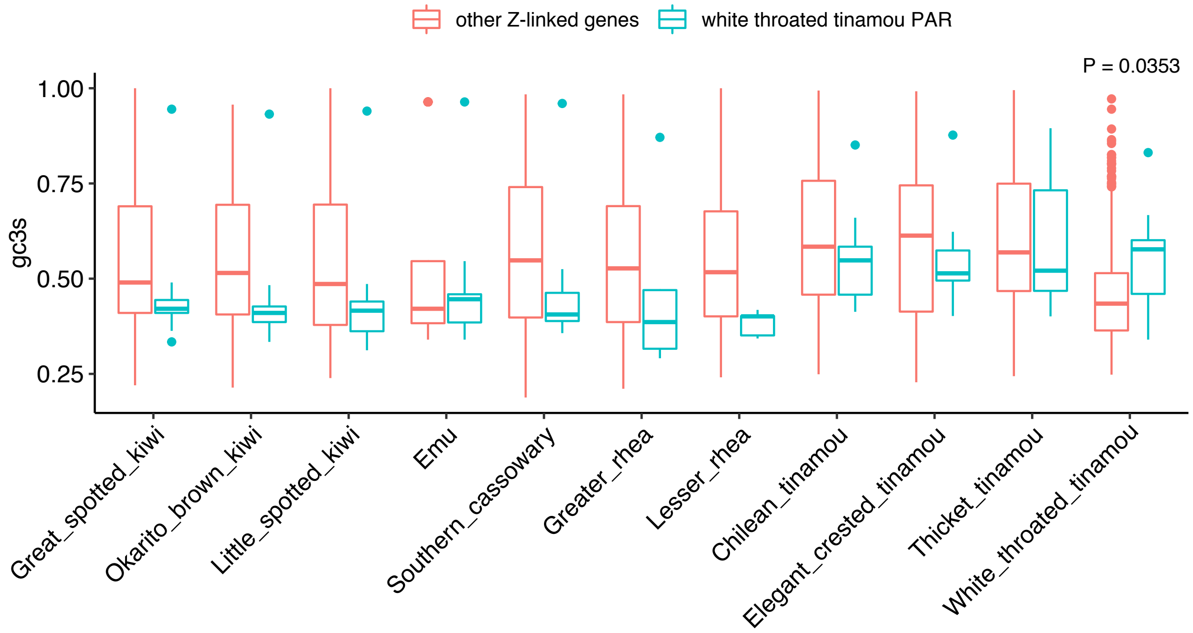


**FIG. S9. The PAR-linked genes in white throated tinamous is GC-biased.** The boxplots show the median of gc3s of nine PAR-linked (genBank ID 104571644, 104571645, 104571646, 104571647, 104571642, 104571648, 104571649, 104571643 and 104571650) gene and the rest Z-linked genes. Their homologous genes in other species are also shown for comparison. The PAR-linked genes of white throated tinamou are GC-biased only in white throated tinamou (P = 0.0353, Wilcoxon rank-sum test).

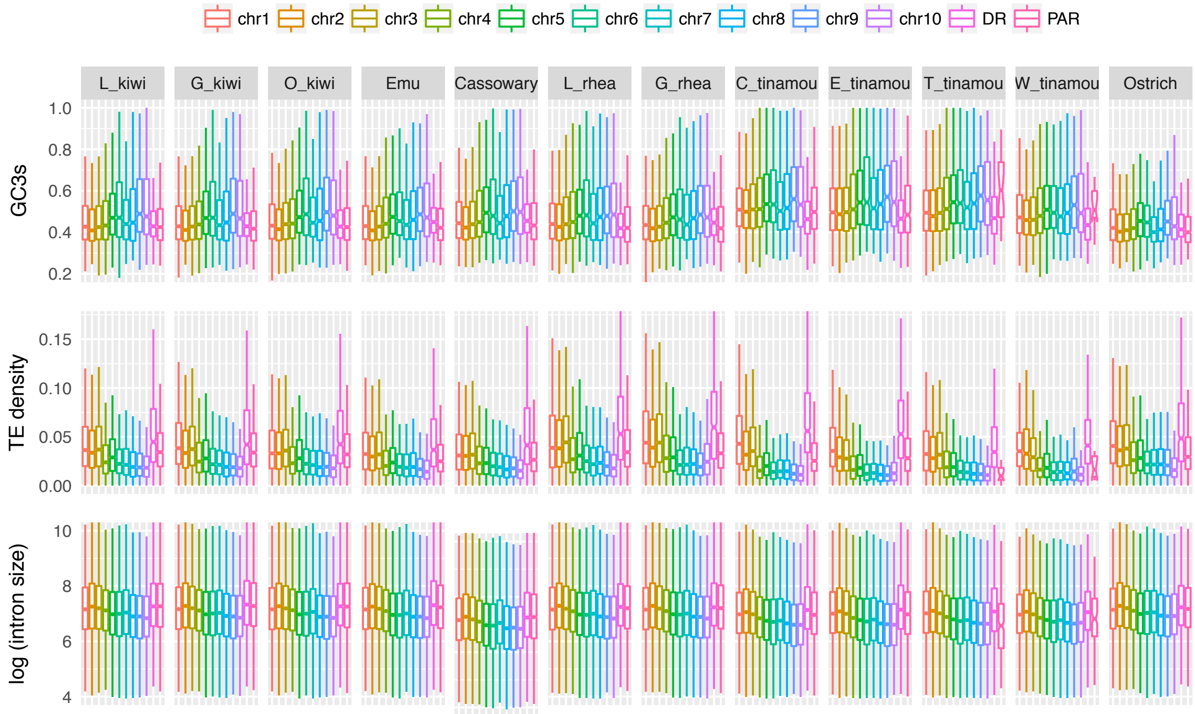


**FIG. S10. Comparison of genomic feature among macro-chromosomes**. GC3s (GC content of synonymous site of the third codon) and exon density show negative correlation with chromosome size, while TE (transposable element) density and intron size show positive correlation with chromosome size.

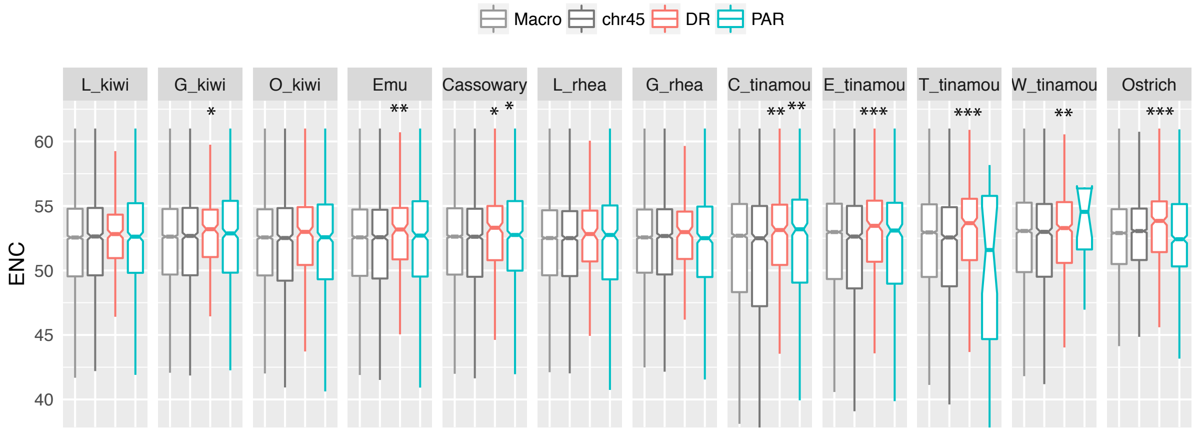


**FIG. S11. Comparison of ENC between PAR/DR and autosomes**. The ENC (Effective Number of Codons) values are higher in DRs for many species, but only for cassowary and Chilean tinamou ENC values are higher in PAR than for autosomes. Asterisks indicate the significant levels of PAR/DR *vs.* chr4/5 comparison (Wilcoxon sum rank test), * <0.05, ** <0.01, *** <0.001.

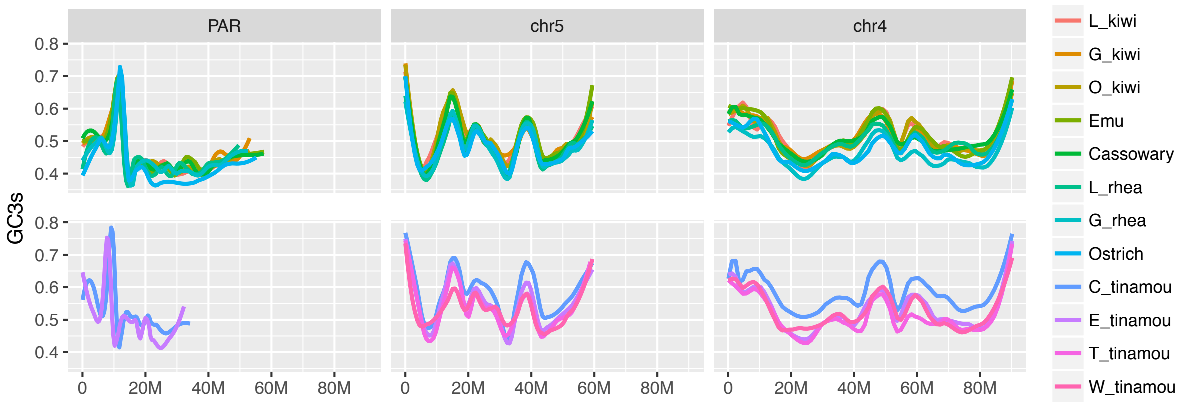


**FIG. S12. Reduced GC3s on the PARs compared to chr5 and chr4**. The location of the PAR-linked genes is based on the pseudo-chromosome Z, and the location of genes of chr4 and chr5 are based on the homologous genes of the chicken genomes. The abbreviation for species names is the same as in FIG S8.

**Table S1. The length of pseudoautosomal region (PAR) and differentiated region (DR) in palaeognaths and selected neognaths**

| Species | PAR Length (bp) | PAR #gene | DR Length (bp) | DR #gene | Reference |
|---|---|---|---|---|---|
| Little spotted kiwi | 53,648,137 | 644 | 27,858,477 | 315 | This study |
| Great spotted kiwi | 53,103,935 | 639 | 27,917,301 | 295 | This study |
| Okarito brown kiwi | 53,052,411 | 655 | 29,311,255 | 345 | This study |
| North Island brown kiwi | ~20M | - | ~65M | - | This study |
| Emu | 59,302,072 | 695 | 21,929,632 | 234 | This study |
| Southern cassowary | 59,264,808 | 749 | 22,782,997 | 295 | This study |
| Lesser rhea | 54,869,205 | 508 | 26,064,611 | 207 | This study |
| Great rhea | 52,553,322 | 604 | 29,742,736 | 214 | This study |
| Chilean tinamou | 34,050,901 | 485 | 36,483,903 | 380 | This study |
| Elegant created tinamou | 32,217,551 | 419 | 33,168,615 | 350 | This study |
| Thicket tinamou | 250,000 | 10 | 71,263,047 | 831x | This study |
| White-throated tinamou | 685,144 | 14 | 62,454,206 | 736 | (Zhou et al. 2014) |
| Ostrich | 52,483,918 | 704 | 31,782,146 | 391 | (Zhou et al. 2014), this study |
| Collared flycatcher | 630,000 | 17 | 68,355,977 | 591 | (Smeds et al. 2014) |
| Zebra finch | 450,000 | 16 | 75,826,118 | 653 | (Singhal et al. 2015) |
| Chicken | 10,000 | 0 | 82,519,921 | 826 | (Bellott et al. 2017) |
| Pekin duck | 1,050,000 | - | 76,500,000 | - | (Zhou et al. 2014) |

Large PAR species are shaded in gray

**Table S2. P-values of Fisher's exact test for overrepresentation of sex-biased on the Z chromosome and PAR.**

| Tissue | Z chromosome Male biased | Z chromosome Female biased | PAR Male biased | PAR Female biased |
|---|---|---|---|---|
| Spleen | **1.26E-23** | 0.062 | 0.303 | 0.122 |
| Gonad | **2.72E-07** | 1.000 | 1.000 | 1.000 |
| Brain | **0.00848** | 0.211 | 0.637 | 0.910 |
| Embryo day15 | **1.52E-05** | 1.000 | 0.214 | 0.747 |
| Embryo day42 | **0.00676** | 0.524 | 1.000 | 0.174 |

**Table S3.** Correlation between chromosome sizes and genomic features

| Species | | GC3s | TE density | Intron size | Exon density | ENC | Intergenic size |
|---|---|---|---|---|---|---|---|
| L_kiwi | r | -0.86 | 0.90 | 0.83 | -0.68 | 0.79 | 0.53 |
| | p-value | 0.00143 | 0.00033 | 0.0033 | 0.03091 | 0.00633 | 0.11574 |
| G_kiwi | r | -0.86 | 0.93 | 0.81 | -0.75 | 0.74 | 0.50 |
| | p-value | 0.00157 | 0.00009 | 0.00434 | 0.01307 | 0.01534 | 0.1394 |
| O_kiwi | r | -0.86 | 0.90 | 0.74 | -0.63 | 0.87 | 0.19 |
| | p-value | 0.00134 | 0.00045 | 0.01365 | 0.0532 | 0.00117 | 0.60513 |
| Emu | r | -0.86 | 0.92 | 0.86 | -0.71 | 0.90 | 0.71 |
| | p-value | 0.00158 | 0.00019 | 0.00125 | 0.02259 | 0.00035 | 0.02122 |
| Cassowary | r | -0.87 | 0.92 | 0.84 | -0.84 | 0.87 | 0.76 |
| | p-value | 0.00105 | 0.00013 | 0.00256 | 0.00209 | 0.00119 | 0.01071 |
| L_rhea | r | -0.88 | 0.90 | 0.87 | -0.77 | 0.77 | 0.62 |
| | p-value | 0.00069 | 0.00039 | 0.0011 | 0.00957 | 0.00951 | 0.05399 |
| G_rhea | r | -0.91 | 0.92 | 0.82 | -0.81 | 0.77 | 0.86 |
| | p-value | 0.00021 | 0.00017 | 0.00406 | 0.00439 | 0.00893 | 0.00124 |
| C_tinamou | r | -0.79 | 0.95 | 0.89 | -0.86 | 0.91 | 0.81 |
| | p-value | 0.00681 | 0.00003 | 0.00059 | 0.00131 | 0.00022 | 0.00447 |
| E_tinamou | r | -0.90 | 0.94 | 0.91 | -0.79 | 0.94 | 0.74 |
| | p-value | 0.00039 | 0.00006 | 0.00022 | 0.00681 | 0.00006 | 0.01391 |
| T_tinamou | r | -0.89 | 0.97 | 0.91 | -0.85 | 0.88 | 0.78 |
| | p-value | 0.00066 | 0 | 0.00023 | 0.0017 | 0.00068 | 0.00735 |
| W_tinamou | r | -0.82 | 0.94 | 0.87 | -0.80 | 0.80 | 0.71 |
| | p-value | 0.00396 | 0.00006 | 0.00119 | 0.00513 | 0.00555 | 0.02226 |
| Ostrich | r | -0.54 | 0.95 | 0.75 | -0.77 | 0.51 | 0.75 |
| | p-value | 0.10623 | 0.00004 | 0.01255 | 0.00933 | 0.1314 | 0.01212 |

**Table S4. Location of Janes et al 2009 BAC sequence in the emu genome assembly.**

| GENBANK RECORD | CHROMOSOME (JANES ET AL 2009) | EMU GENOME LOCATION | EMU GENOME CHROMOSOME |
|---|---|---|---|
| EU200931 | Autosome | not determined | not determined |
| EU200931 | Autosome | not determined | not determined |
| ET041500 | Autosome | not determined | not determined |
| ET041501 | Autosome | not determined | not determined |
| ET041502 | Autosome | not determined | not determined |
| ET041515 | Autosome | not determined | not determined |
| ET041512 | Autosome | not determined | not determined |
| ET041513 | Autosome | not determined | not determined |
| AB002056 | PAR | presumed assembly gap | Z (PAR) (1) |
| AB006694 | PAR | scaffold_221: 152067-154756 | Z (DR) |
| AY095498 | PAR | scaffold_13:6582073-6583283 | Z (DR) |
| AB006695 | PAR | scaffold_239:1028675-1030884 | Z (PAR) |
| ET041507 | PAR | scaffold_16:5843173-5843899 | chr5 |
| ET041520 | PAR | scaffold_66:2150084-2282118 | chr7 |
| ET041521 | PAR | scaffold_66:2150084-2282118 | chr7 |
| ET041516 | PAR | scaffold_14:4429102-4300170 | chr4 |
| ET041517 | PAR | scaffold_14:4429102-4300170 | chr4 |
| ET041508 | PAR | scaffold_19: 1980997-2079372 | Z (DR) |
| ET041509 | PAR | scaffold_19: 1980997-2079372 | Z (DR) |
| ET041510 | PAR | scaffold_19: 1980997-2079372 | Z (DR) |
| ET041518 | PAR | scaffold_106:822019-944625 | chr8 |
| ET041519 | PAR | scaffold_106:822019-944625 | chr8 |

(1) determined by alignment to other palaeognaths
